# Collective Evolution Learning Model for Vision-Based Collective Motion with Collision Avoidance

**DOI:** 10.1101/2022.06.09.495429

**Authors:** David L. Krongauz, Teddy Lazebnik

**Author notes:** These authors contributed equally to this work.

## Abstract

Collective motion (CM) takes many forms in nature; schools of fish, flocks of birds, and swarms of locusts to name a few. Commonly, during CM the individuals of the group avoid collisions. These CM and collision avoidance (CA) behaviors are based on input from the environment such as smell, air pressure, and vision, all of which are processed by the individual and defined action. In this work, a novel vision-based CM with CA model (i.e., VCMCA) simulating the collective evolution learning process is proposed. In this setting, a learning agent obtains a visual signal about its environment, and throughout trial-and-error over multiple attempts, the individual learns to perform a local CM with CA which emerges into a global CM with CA dynamics. The proposed algorithm was evaluated in the case of locusts’ swarms, showing the evolution of these behaviors in a swarm from the learning process of the individual in the swarm. Thus, this work proposes a biologically-inspired learning process to obtain multi-agent multi-objective dynamics.

**Author summary:** Multi-agent multi-objective tasks are common in nature with examples as collective movement in birds and economic management in humans. These problems are famous for being convoluted to efficiently solve. Nonetheless, nature has been successfully solving it for millennials using an evolution strategy. A prominent example is a task of flocking performed by multiple species, which involves both collective motion and collision avoidance. In our work, we simulate agents that are able to learn behaviors on the individual level, that in turn translate into the desired group (multi-agent) behavior. Using nature-inspired genetic algorithms and reinforcement-learning methods, the agents are successfully implementing a collective behavior similar to the one encountered in nature.

## 1 Introduction

Collective motion (CM) is a phenomenon occurring in homogeneous populations of interacting individuals [1–4]. It is manifested in multiple forms in nature such as locusts aggregate into groups [5, 6], birds flocking in an organized manner [7, 8], and fish schools responding to predators and showing migratory movements [9, 10]. During the last few decades, researchers seek to elucidate the mechanisms allowing animals to achieve synchronized motion while lacking centralized control. Empirical data from experimental observations [11, 12] is analyzed with the aim to uncover individual’s behavior using mathematical and computational models [13–15].

A large body of work can be categorized as agent-based models, where the collective dynamics are explained by the action on the individual-level [16]. One of the leading approaches in agent-based simulation for CM is Self-Propelled-Particles, popularized by the model proposed by Vicsek et al. [17], where agents align their velocity according to short-range interaction with neighboring agents, commonly known as the metric-based approach. In this group of models [18–20], agents are moving with constant speed [17], aligning their velocity direction in response to inter-agent interactions. Moreover, a notable subgroup of models is the Avoidance-Attraction-Alignment group [5, 21, 22] where agents are governed by three types of superimposed forces: aligning force (For example see [17]), separating force for collision avoidance, and cohesion to the center of mass of the group. Couzin et al. [23] showed how efficiently the information can transfer between members of animal groups. Another model incorporating metric-based interaction proposed by Bhattacharya and Vicsek [24], described the processes involved in birds flocking and synchronized landing. In addition, describing the dynamics and synchronization of general clusters of self-propelling-particles utilize the metric approach [25, 26].

Contrary to the metric-based approach, the *topological-based* approach has suggested that birds flock according to interactions with a fixed number of neighbors regardless of their distance from each other [27]. Namely, this approach differs from the metric in neighbors’ capacity, since theoretically the entire swarm can be classified as ’neighbors’ if they are located within a predefined distance from each other. Shang and Bouffanais [28] proposed a method to find the optimal number of neighbors for the topological-based approach while taking into consideration the interacting self-propelled particles that aim to reach a consensus. Camaperi et al. [29] compared the metric-based and topological-based approaches, and showed superior stability for the latter due to the fact that particles displaying a more uniform distribution lead to higher cluster stability. Kumar and De [30] explored and compared the flocking efficiency of metric-based and topological-based approaches.

Although these models reproduce aspects of the macro-dynamics of natural swarms [31, 32], they do not address the problem of information acquisition by the agents, since models relying on globally available information. For instance, accurate positions and velocities of agents in the environment are commonly assumed to be fully known to the focal agent [17, 18, 21]. Nonetheless, in reality, agents’ perception of the environment is mediated by their sensory systems (e.g. vision, hearing, echolocation, magnetic field sensing etc.) [33–36]. Distinctively, objects in the environment are reflected on the retina of the focal agent’s eye and later translated to the brain. This input is used as part of the agent’s decision-making process, allowing an individual to act upon the visual stimuli it observed [37, 38]. Indeed, many animals use vision as the main sense for coordination and navigation [39–41]. Hence, to obtain more biologically accurate modeling of collective motion, agents in the swarm are assumed to estimate an object’s location and velocity using vision-based sensing.

Multiple models that encompass visual projection in CM modelling were proposed [42–45]. Strandburg-Peshkin et al. [44] explored visual sensor networks and their role in information transfer between schools of fish. Bastien and Romanczuk [46] used elementary occlusions of visual rays to create avoidance and self-organizations in disc-like shaped agents. The authors demonstrated several group behaviors with steering by angular visual occlusions. A follow-up work introduced an additional collision avoidance and demonstrated collective behavior in the presence of static obstacles in several settings [47]. Both models are oblivious to object recognition, thus any agent-specific features are ignored. In the present work, we propose and demonstrate a different approach in which the shape and size of the agents in the swarm are effective the decision-making process due to the vision-based input.

Several attempts to model CM using learning models are presented [48–50]. Durve et al. [48] developed both Vicsek-inspired CM teacher-based reinforcement learning and concurrent learning. We conclude that the velocity alignment mechanism allows spontaneously emerge from the minimization of neighbor-loss (e.g., agents that are not in the visual radius between two observations) rate. Thus, the model learn to keep group cohesion when the perception is limited to the velocity of neighboring agents [48]. They assume the focal agent is accurately aware of the location and velocities of the neighboring agents. Into the bargain, Young and La [49] presented a hybrid system that achieves an efficient CM behavior for escaping attacking predators while maintaining a flocking formation using multiple reinforcement learning methods. Additionally, it was shown that independent learning with and without function approximation proved to be unreliable in learning to flock together towards the same target. Similarly, in our work, it is assumed the focal agent is accurately aware of the location and velocities of the neighboring agents while solving a multi-objective task. Nonetheless, they were not able to obtain a collective behavior by learning a policy at the individual level as obtained by Durve et al. [48]. Lopez-Incera et al. [50] studied the collective behavior of artificial learning agents driven by reinforcement learning action-making process and with abstract sensing mechanism, that arises as they attempt to survive in foraging environments. They design multiple foraging scenarios in one-dimensional worlds in which the resources are either near or far from the region where agents are initialized and show the emergence of CM [50]. While their work took into consideration a sensing-based approach, they investigated a single objective behavior in an extremely simplistic scenario.

The novelty of the proposed work lies in the development of a Collective Evolution Learning (CEL) algorithm that allows learning a multi-objective collective behavior based on the optimization of a single member’s decision-making process even with noisy input data. In particular, we show that CEL is able to achieve flocking with inter-agent collision avoidance using only vision-based inputs.

Formally, our model produces vision-based collective motion (VCM) group behavior combined with collision avoidance (CA) or VCMCA for short. The model is obtained by simulating the evolution process of an animal population that learns the desired behavior over time using a trial-and-error mechanism. We evaluate the performance of the algorithm to obtain a VCMCA model in different settings, showing robust results. The remaining of the manuscript’s structure is shown in Fig. 1.

**Fig 1.**
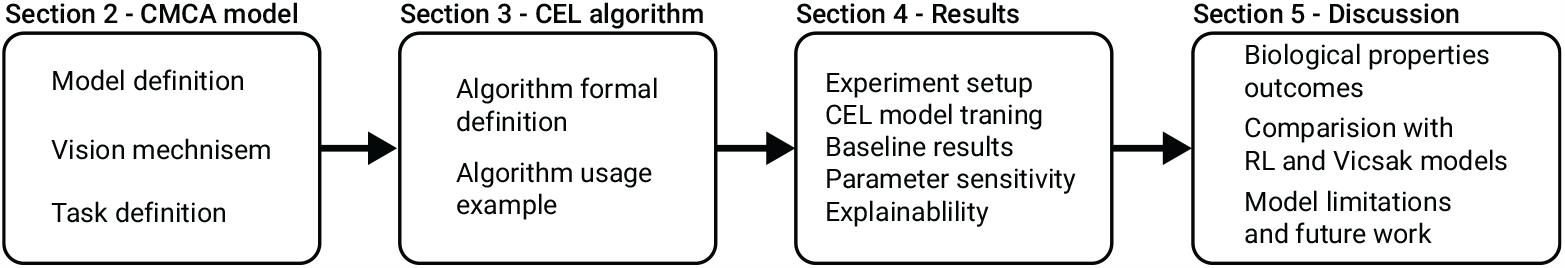
A schematic view of the manuscript’s structure. Section 2, outlines the mathematical formalization for the CM with CA in a two-dimensional setting with agents that based their decisions on a vision mechanism. In Section 3, a novel computational model for vision-based CM with CA, called VCMCA, is proposed. In Section 4, the performance of the VCMCA using several *in silico* experiments is evaluated. In Section 5, biological outcomes, comparison with other RL- and Viscak- based models, and the limitations of the model are discussed.

## 2 Vision-Based Collective Motion with Collision Avoidance

In this section, we describe the proposed VCMCA model, constructed from three components: the vision-based input mechanism on the agent level, the movement and collision physical mechanism, and the swarm-level CM and CA dynamics. A table that summarizes the parameters used in the model, their definition, and notations are provided in the Appendix.

### 2.1 Settings and vision-based input

In order to represent the open-field conditions that characterize swarming organisms’ environments in a simple manner, we define a two-dimensional world with two-dimensional agents, in a similar fashion to Vicsek’s model [24]. We use the metric-based approach to define the neighbors of an agent.

The model (*M*) is defined by the tuple *M* := (*E, P*) where *E ⊂* ℝ^2^ is the environment and *P* is a fixed sized (‖*P*‖ := *N*) population of rectangular-shaped agents populating it. Formally, the environment, *E*, is a continuous space with height *H* ∈ ℕ and width *W* ∈ ℕ with a toroidal geometrical configuration. Namely, a two-dimensional rectangle with periodic boundary conditions. Each agent, *p* ∈ *P*, is defined by a tuple *p* := (*l, v, a, h, w, r, γ*) where *l* = (*l*_*x*_, *l*_*y*_) ∈ *E* is the location of the agent’s center of mass, *v* ∈ ℝ^2^ is the current velocity vector of the agent, *a* ∈ ℝ^2^ is the current acceleration vector of the agent, *h, w* are the agent’s height and width, respectively, and *r* is the metric-based sensing radius of the agent.

The parameters used for the agent’s visual mechanism are inspired by previous works dealing with vision-based information acquisition of flocking animals [45, 51]. Formally, the vision-based input is parameterized by *γ* := (*ψ*, 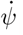, *θ*, 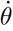), where *θ* and *ψ* are the angular position and area subtended on the retina of the focal agent, respectively, and 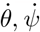, are their respective derivatives in time.

Based on the sensing radius, *r*, we define the *local environment* of a focal agent *p*_*o*_ ∈ *P* (we use *o* to denote parameters related to the focal agent) to be the subset of agents *P*^∗^ that are located in a sensing distance from the focal agent *P*^∗^ := { *p*_*i*_ ‖*P* : ‖*l*_*i*_ *l*_*o*_ ∈< *r*_*o*_ } such that *i* ≠ *o*. Now, each agent in the local environment (*p*_*i*_ ∈ *P*^∗^) of a focal agent *p*_*o*_ is sensed using the vision mechanism if it is not fully obscured by other agents. The vision-based properties (*γ*) are computed for each neighbor *n* in the *local environment* of the focal agent, resulted in a vector *ν* ∈ ℝ^|*γ*|·|*P*^ ∗|, such that *ν* := [*γ*_1_, …, *γ*_|*P*_ ∗|]. The subtended angular area *ψ* for each neighbor *n* is obtained as follows:

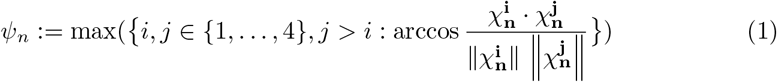

where 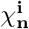 are the relative positions of the corners of n-th neighbor as seen in the focal agent frame. Formally, the lab frame coordinates of the corners of the nth neighbor:

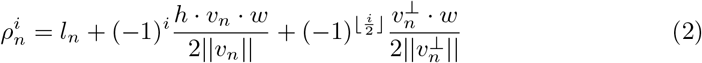

such that index *i* ∈ *{*1, 2, 3, 4*}*, 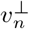 is the perpendicular vector to the velocity vector *v* of the neighbor agent, and *l*_*n*_ is the position of the *n*th neighbor’s center of mass. The coordinates in the focal agent (o) frame would result in 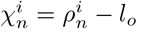. Thus, the angular position parameter *θ* of the *n*th neighbor is defined as follows:

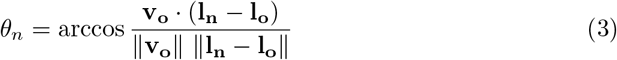

An agent can be fully latent to the focal agent by other agents in its local environment. In such a case, it is not taken into consideration in the vision-based computation. A schematic view of the focal’s agent vision-based sensing (*γ*) is shown in Fig. 2. More details about the computation of Eqs. (2) and (3) are provided in the Appendix.

**Fig 2.**
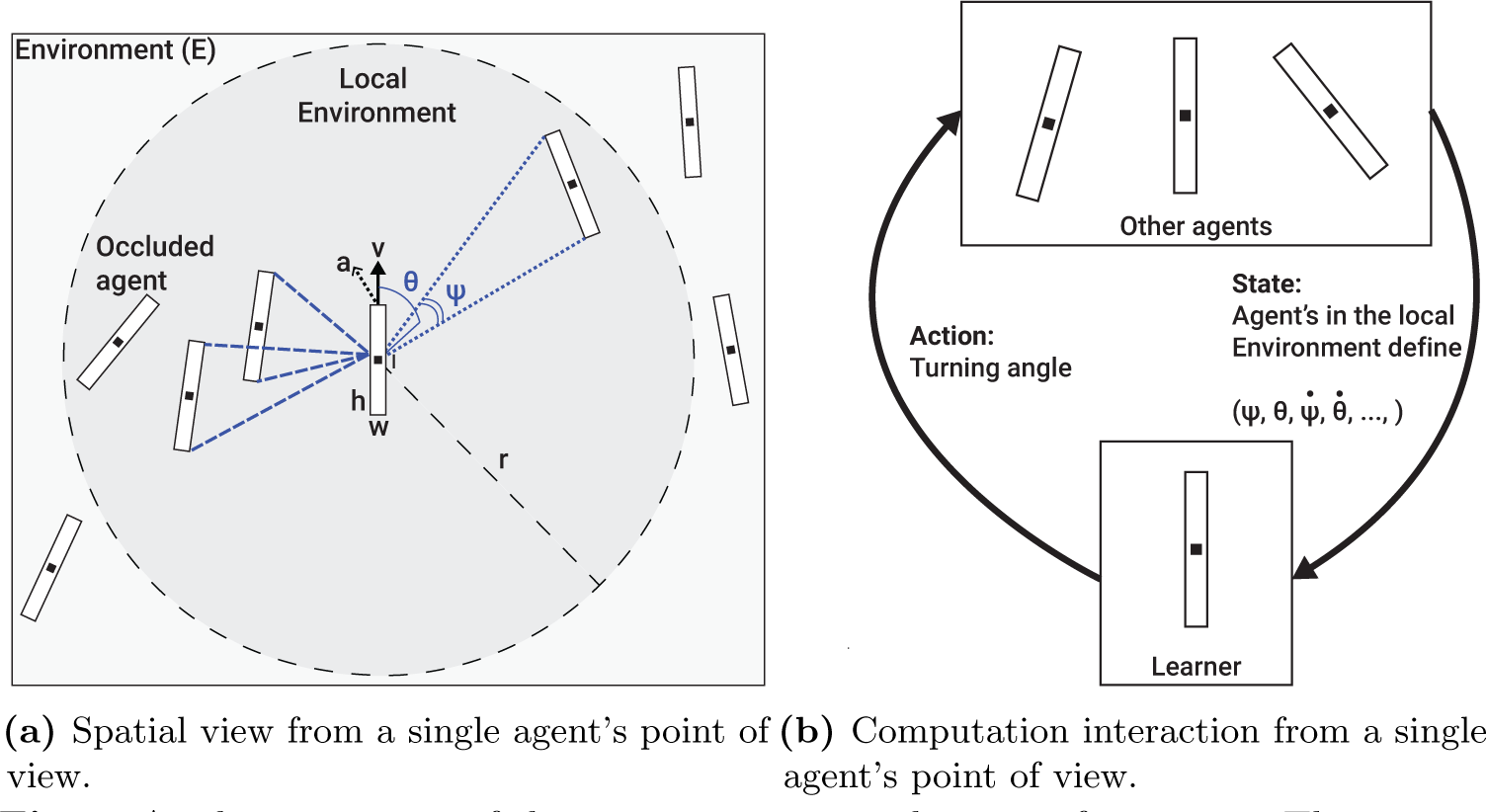
A schematic view of the vision sensing mechanism of an agent. The computation is presented for a single focal agent but performed symmetrically for all the agents in the population.

### 2.2 Agent’s movement and collision dynamics

The proposed model has a discrete, synchronized clock that all the agents, *P*, follow. At the beginning of the simulation (*t*_0_), the locations and velocities of the agents’ population (*P*) are chosen at random from a pre-defined distribution. Then, for each clock tick (marked by *t*_*i*_ for the *i*_*th*_ tick) the following happens: first, each agent *p*_*o*_ ∈ *P* senses its local environment and computes its action *x*(*t*) ∈ ℝ^2^ based on a given decision-making model, in a random order. Second, each agent moves as follows:

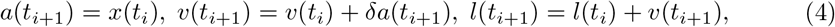

where *δ* ∈ *ℝ*^+^ is a weighted variable indicates the influence of the desired direction (*x*(*t*_*i*_)) on the current direction. Due to the toroidal geometrical configuration of the environment, if *l ∉ E* then 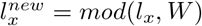 and 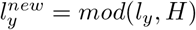.

In case two agents *p*_1_ ∈ *P, p*_2_ ∈ *P* collide, a collision protocol is employed: first, the simulation identifies the both colliding agents to be either *active* or *passive* colliders. Namely, in a head-tail collision, the head colliding agent is regarded as the active and the tail colliding as the passive. In the case of a frontal collision, both parties are regarded as active colliders. The active colliding agent(s) are stopped (i.e., *v*(*t*_*i*_) = 0) while the passive colliding agent’s velocity follows purely-elastic collision:

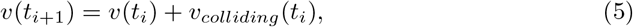

where *v*_*colliding*_(*t*_*i*_) is the velocity of the colliding agent at the time of the collision (*t*_*i*_). A schematic view of the perception-action-making loop of a single agent is presented in Fig. 2b.

### 2.3 Collective motion and collision avoidance

We define the agents’ population CM metric by the degree of alignment of the velocity headings of the simulated agents, in a similar fashion to other CM models’ metrics [5, 17]. Formally, the CM metric is a function *F* :ℙ → ℝ where𝕡 is the space of all possible agent populations. *F* (*P*) is defined as follows:

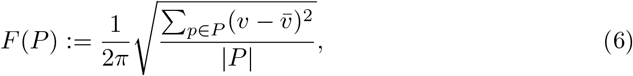

where 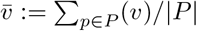.

In addition, we define a collision between two agents *p*_1_ ∈ *P* and *p*_2_ ∈ *P* taking place if and only if:

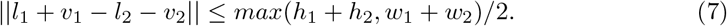

The motivation behind this definition is to check if two agents *p*1, *p*2 would collide after they both make a move. In order to evaluate if a collision happened, we check if the distance between the agents’ centers of masses (*l*_1_ + *v*_1_ and *l*_2_ + *v*_2_) is smaller than their width or height. This definition is optimal when assuming circular-shaped agents. Nonetheless, for non-circular-shaped agents, such as in our case, is considered a bit aggravating as there are angles in which the condition is met but the agents do not physically collide. Therefore, a function *C* : ℙ → ℕ such that *C*(*P*) counts the rate of collisions between the agents of the population to the possible number of collisions in a single point of time, is used as the CA metric. In our context, *C* is normalized to [0, 1] by dividing *C* by the largest hypothetical number of collisions possible in a single step in time.

Based on these two metrics, the objective of the CM with CA takes the form:

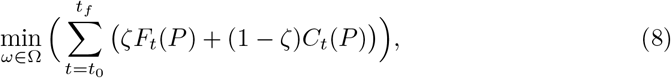

where Ω is the space of all possible policies of the agents and *ζ* ∈ [0, 1] is the weight of the CM score compared to the CA score. The optimization is done on any finite duration in time [*t*_0_, …, *t*_*f*_ ] ⊂ ℕ such that *t*_0_ < *t*_*f*_ and *F*_*t*_ and *C*_*t*_ represent the CM and CA at time *t*, respectively.

## 3 Collective Evolution Learning Algorithm

### 3.1 Motivation

The CM behavior of locusts is believed to develop over time throughout an evolution process. The locusts try to adapt to their environment and generate the next population of locusts. During this process, multiple locusts experienced similar situations and operate slightly differently based on unique properties each locust has which resulted in different outcomes. Locusts that perform better were more likely to reproduce and keep their behavior in the population over time. As such, the wanted behavior (e.g., CM with CA) emerged throughout the collective experience of the locust and improved over time using a natural trial-and-error approach.

Based on this motivation, we propose a reinforcement learning (RL) based algorithm (*A*) called Collective Evolution Learning (CEL for short). The proposed algorithm is based on the assumption the agents in a swarm are identical and share common knowledge (policy). Intuitively, the CEL algorithm receives the observable parameters (*γ*) of a subset of agents (*P*^∗^ *⊂ P/{p}*) in the local environment (*R*) of an agent (*p*) and returns the action (*x*) that best approximate the optimal action of the CM with CA objective metric (Eq. (8)) using a Q-learning based model [52, 53]. Since both CM and CA are swarm-level metrics, CEL has an intermediate genetic algorithm (GA) based model [54–58] to optimize CM and CA on the agent’s level. Moreover, to obtain a well-performing Q-learning-based model, one is required to well sample the state space of the model [59]. In order to do that, CEL introduces a layer on top of the Q-learning model that measures its performance for different scenarios and uses the K-nearest neighbors (KNN) algorithm [60] to define a new sampling strategy for further training of the Q-learning model. A schematic view of the model training, validation, and testing phases with the interaction between them is presented in Fig. 3.

**Fig 3.**
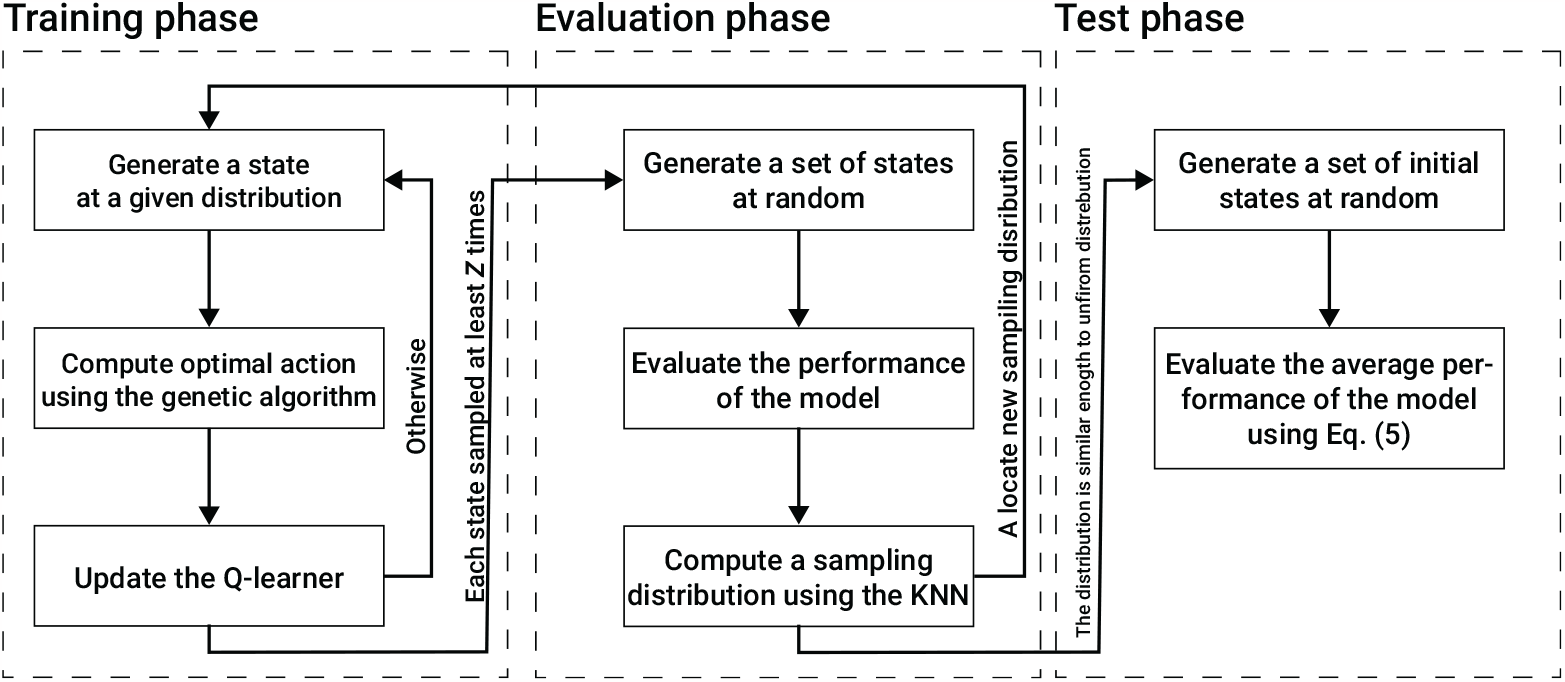
A schematic view of the model training, validation, and testing phases with the interaction between them.

### 3.2 Swarm dynamics using a Q-learner

Formally, CEL is a function *A* : ℕ^4*k*^ → ℕ such that *A*(*ν*) → [*x*_1_, …, *x*_*n*_] where *k* ∈ ℕ is the maximum possible number of agents that an agent *p* ∈ *P* can sense in a single point in time multiplied by the number of vision-based parameters per agent (i.e., *ψ, θ*, 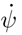, and 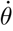), *ν* ∈ℝ^4*k*^ is the outcome of the vision-based sensing of an agent *p* ∈ *P*, and 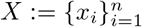 is the set of all possible actions an agent can make. Following this definition, one can treat *A* as a policy function getting a state *ν* for each agent and return the action *x* it should perform. To learn the policy *A*, we use a Q-learning-based model. The Q-learner stochastically maps each state *ν* to the set of possibles actions 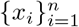 by assuming a discretization of both the input and output spaces. Hence, it is assumed that the number of possible actions is finite and that *ν* ∈ ℕ ^4*k*^ is obtained by the discretization of each vision parameter to define a classification problem rather than a regression problem.

The process of obtaining the model, is divided into three main phases: training, evaluation, and testing. The training and evaluation phases are executed sequentially multiple times until a stop condition is satisfied. Afterward, the testing phase is taking place to compute the VCMCA score of the obtained model.

The training phase is further divided into the first time the training phase is utilized and any other time afterward. For the first time, the Q-learning matrix is initialized with all possible states and their actions such that each action has a uniform distribution to be chosen for each state. Afterward, and for each utilization of the training phase, the training phase includes three linear processes. First, a scenario including zero, one or more (limited by *k*) neighbor agents, and a focal agent is generated, at random and in an iterative manner. Based on this scenario, the Q-learner’s state is computed from the focal agent’s point of view. Second, an approximation to the optimal actions for this scenario is obtained using a GA-based model aiming to optimize the CMCA for *η* ∈ ℕ steps in time. This step is described in detail in Section 3.3. Lastly, the results of the algorithm are used to update the Q-learner. The Q-learner is updated by adding to the current value a score obtained by the GA process multiplied by some pre-defined learning rate, *λ* ∈ [0, 1]. This process repeats itself for *Z*_*train*_ ∈ ℕ times.

The evaluation phase includes three linear processes. First, a set of scenarios of size *Z*_*validate*_ ∈ ℕ are generated at random. For each scenario, the Q-learner’s state is computed from the focal agent’s point of view. Pending, the performance of the Q-learner for each state is computed by simulating *η* steps in time such that both the focal and neighbor agents follow the Q-learner’s model. Afterward, CEL uniformly samples the state space and computes the estimated performance of the model on these values using the distance-weighted KNN algorithm with *y* ∈ ℕ neighbor data points. The outcome of the last step is a 4*k*-dimensional distribution function. If the total variation distance between the obtained distribution and a uniform distribution is larger than a pre-defined threshold *κ* ∈ ℝ^+^ then CEL gets back to the training phase with the obtained distribution to sample from for the next set of training samples. Otherwise, CEL moves to the testing phase.

The testing phase includes three linear processes. First, CEL generates a set of simulation scenarios of size *Z*_*test*_ ∈ ∕, at random. Notably, the simulation scenarios are of the entire population and not of a single focal agent and its neighbor agents as used during the training and evaluation phases. Second, for each simulation scenario, the simulation is performed for 1 << *T* ∈ℕ steps in time such that all the agents in the population follow the Q-learner model. Lastly, the performance of the CEL model is defined to be the average CMCA score of the simulations.

### 3.3 Agent’s decision making using a genetic algorithm

An approximation to the optimal actions for a given scenario is obtained using a GA-based model aiming to optimize the CMCA score of a single agent over a short duration of time. To use the GA, we first define a local, individual objective function, based on the collective metric Eq. (8), as follows:

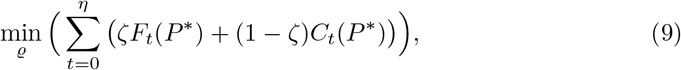

where *P*^∗^ *⊂ P* is the subset of agents in the local environment of a focal agent *p* ∈ *P*, and *ϱ* is a gene used by the GA. This definition assumes a semi-Markovian property [61] to the action-making processes of the agents which means that the agents are aiming to perform the optimal action for the next state, taking into consideration the current and previous states.

First, initialization of the gene population is taken place. Formally, a gene’s population of size *τ* ∈ ℕ such that each gene (*ϱ*) is defined by a vector of size *η* corresponding to the number of steps in time the focal agent needs to perform is generated. Each value of the vector is an action *x* ∈ *X*. Afterward, for *g* ∈ *ℕ* generation (iterations), three operations took place: cross-over, mutation, and next generation. To be exact, the cross-over operator 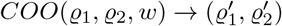 gets two genes *ϱ*_1_, *ϱ*_2_ and a weight vector *w* ∈ [0, 1]^*η*^ and returns two new genes 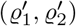 such that 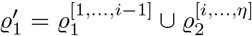 and 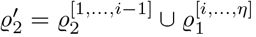, where *i* ∈ [2, *η*] is an index of the genes’ vector sampled in a distribution defined by *w* [62]. *The mutation operator MO*(*ϱ*) → *ϱ*^*′*^ gets a gene *ϱ* and returns an new gene *ϱ*^*′*^ such that one of the values of *ϱ* altered with a different action in a uniformly distributed manner. The next generation operator (NGO) leaves a portion *ρ* ∈ [0, 1] of the genes with the highest fitness score while the remaining genes are chosen by a distribution defined by the *L*_1_ normalization of the fitness scores of the remaining genes [63]. Once the GA finished its calculations, the gene with the best fitness score is taken. From this gene, the first action is provided for the Q-learner.

Notably, one does not have to use the GA approach as in theory the Q-learning algorithm should coverage to a similar policy without the GA component. However, as the number of samples that results in better fitness is really small compared to the action space’s size, a Q-learner would require a very large number of samples to achieve similar results. As such, it would cause a much longer training process.

A time complexity analysis of CEL is provided in the Appendix.

## 4 Results

To evaluate the ability of the CEL algorithm to obtain a VCMCA model and explore the influence of several properties of the agents on the model’s performance, we conduct three experiments. First, measure the performance of the proposed algorithm compared to two other approaches - vision-based CM without CA and random walk. Second, we analyze the influence of population density, agent’s height to width ratio, and sensing radius on the VCMCA model’s performance. Finally, we approximate the VCMCA model obtained by the CEL algorithm using an explainable Random Forest [64] model to extract the rule-base structure for the agent’s action-making process, allowing the user to interpret the connection between the visual stimuli and the reaction of the agent to the visual stimuli.

### 4.1 Setup

We implemented the VCMCA model and CEL algorithm as a simulator for the case of locusts. We chose locusts since, at the beginning of the flocking, they move on the ground, which is approximated by a 2D plan. However, the model can be adapted to other species as well. The simulation is divided into two phases: 1) agent’s individual VCMCA model training using the CEL algorithm; and 2) usage of the obtained VCMCA model in the locust swarm simulation. The default hyperparameters’ values used in the simulation are picked to represent a population of locusts that become locusts. The parameters with their descriptions and values are summarized in Table 1. If not stated otherwise, these hyperparameters’ values are used across all experiments. In particular, the agent’s height to width ratio is set to be 7*/*1. This ratio is obtained by rounding the closest natural number to the average ratio of 50 locusts. To be exact, we manually tag the bounding box of 50 upper-view locust images, obtaining an average height-to-width ratio of 6.78 with 1.48 standard deviation. Specifically, the locust images were of the *Schistocerca cancellata* and *Schistocerca gregaria* species since these are known to exhibit swarming behaviors and density-dependent phase transition [65, 66]. In addition, group population sizes were set with accordance to available biological records of locust swarms [66, 67]. Though the exact density values are one order of magnitude larger than the simulations, we concentrate on the emerging phases of the flock when the density is yet to reach its maximal numbers. The model uses abstract discrete time steps. In order to calibrate the simulation’s abstract time step, we define each time step to be the average duration required by the average locust (agent) to make a decision and move [68].

**Table 1.** The description of the model’s parameters and their default values.

| Parameter definition | Symbol | Value |
| --- | --- | --- |
| Environment's ($E$) width [$m$] | $W$ | 5 |
| Environment's ($E$) height [$m$] | $H$ | 5 |
| Population size [1] | $\vert P\vert $ | 30 |
| Agent's initial velocity [$m/s$] | $v$ | $(U[0, 0.15], U[0, 0.15])$ |
| Agent's initial acceleration [$m/s^2$] | $a$ | (0, 0) |
| Agent's height [$m$] | $h$ | 0.07 |
| Agent's width [$m$] | $w$ | 0.01 |
| Agent's action influence [1] | $\delta$ | 0.05 |
| Agent's sensing radius [$m$] | $r$ | 0.21 |
| CM to CA weight [1] | $\zeta$ | 0.5 |
| The number of steps in time for a single simulation [1] | $T$ | 500 |
| CEL algorithm's maximum number of agents in the local environment [1] | $k$ | 8 |
| CEL algorithm's state (input) space discretization [1] | - | 0.01 |
| CEL algorithm's action (output) space discretization [1] | - | 15 degrees |
| CEL algorithm's training samples [1] | $Z_{train}$ | 10,000,000 |
| CEL algorithm's validation samples [1] | $Z_{validate}$ | 100,000 |
| CEL algorithm's testing samples [1] | $Z_{testing}$ | 450,000 |
| CEL algorithm's validation stop condition parameter [1] | $\kappa$ | 0.05 |
| CEL algorithm's genetic algorithm mutation mean and standard deviation [1] | $\mu, \phi$ | 0.1, 0.05 |
| CEL algorithm's genetic algorithm number of generations [1] | $g$ | 10 |
| CEL algorithm's genetic algorithm population size [1] | $\tau$ | 20 |
| CEL algorithm's genetic algorithm steps in time [1] | $\eta$ | 5 |
| CEL algorithm's KNN's number of neighbors data points [1] | $y$ | 3 |
| CEL algorithm's genetic algorithm royalty rate [1] | $\iota$ | 0.05 |

### 4.2 CEL algorithm training

We train the locusts’ VCMCA model using the CEL algorithm for *n* = 30 times to evaluate the average relationship between the number of training samples and the obtained VCMCA model’s performance due to the stochastic nature of the algorithm. In Fig. 4 the blue line indicates the training fitness’ function value (i.e., Eq. (9)), and the green line indicates the CMCA score (i.e., Eq. (8)). The results are shown as mean ± standard deviation. The vertical dashed-yellow lines indicate the validation phase during the training process as described in Fig. 3.

**Fig 4.**
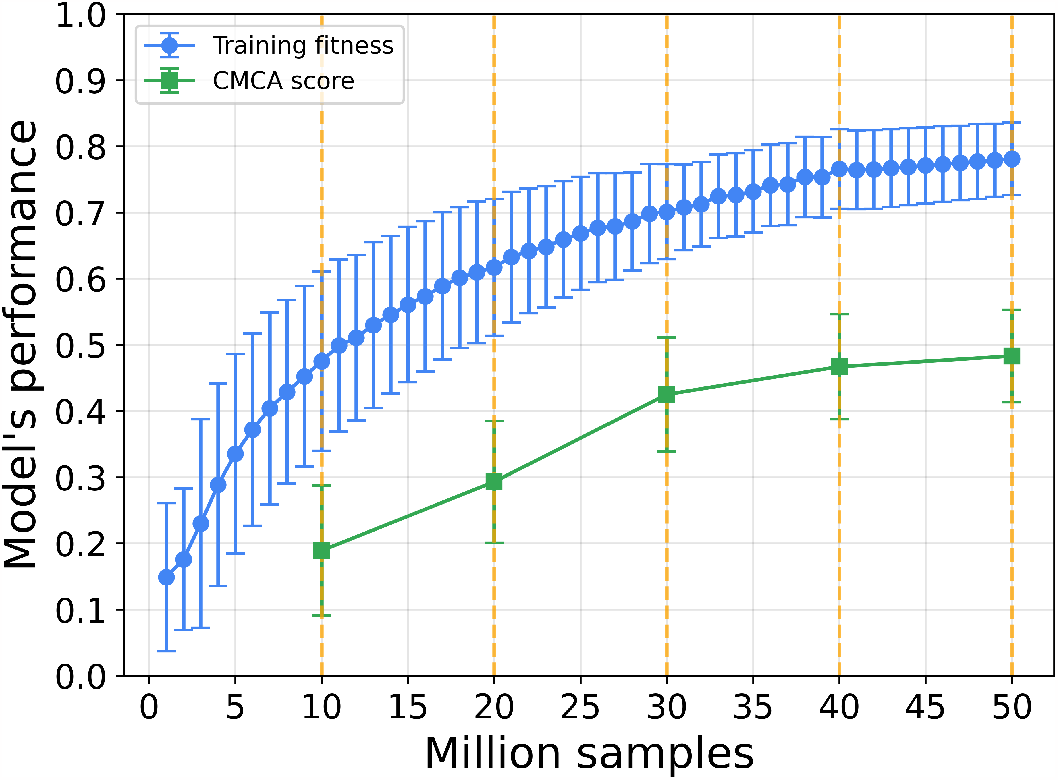
Performance of the learner as training progresses. The training fitness is the agent’s score in the testing phase. The CMCA score is computed by using the trained agent in the simulation. The CMCA score is complement to 1 of the CMCA objective function (Eq. 8). The results are shown as mean ± standard deviation of *n* = 30 repetition. Namely, the error bars are one standard deviation. The vertical dashed-yellow lines indicate the validation phase during the training process. The training is converged after five validations for *κ* = 0.05, *Z*_*training*_ = 10000000, and *Z*_*validate*_ = 100000. **(a)** At the beginning of the simulation (*t* = 0). **(b)** At the end of the simulation (*t* = 500).

We fitted the average training fitness and CMCA score using the least mean square method, obtaining *F*_1_ : *fitness* = 0.189*ln*(*samples*) + 0.053 and *F*_2_ : *cmca* = 0.196*ln*(*samples*) − 0.268 with coefficient of determination *R*^2^ = 0.989 and *R*^2^ = 0.972, respectively. Hence, the functions *F*_1_ and *F*_2_ are well-fitting the dynamics. Since both functions are monotonically increasing, it shows that there is a connection between the CMCA score of an individual and the population’s CMCA score. Similarly, the Pearson correlation between the training fitness and CMCA scores in the validation stages is *R*^2^ = 0.971 with a p-value of 0.0057. Thus, the scores are strongly linearly correlated as well.

Screenshots of the simulator at the beginning and end of the simulation of an average performance VCMCA model after 50 million training samples are shown in Figs. 5a and 5b, respectively. Each agent is represented by a black rectangle with a surrounding gray circle that represents the sensing radius of the agent and its index number. The agents marked in dashed blue indicate that they have collided. One can see that in Fig. 5a that the agents are heading in random directions while in Fig. 5b the agents heading in the same direction (top-right corner), obtaining CM.

**Fig 5.**
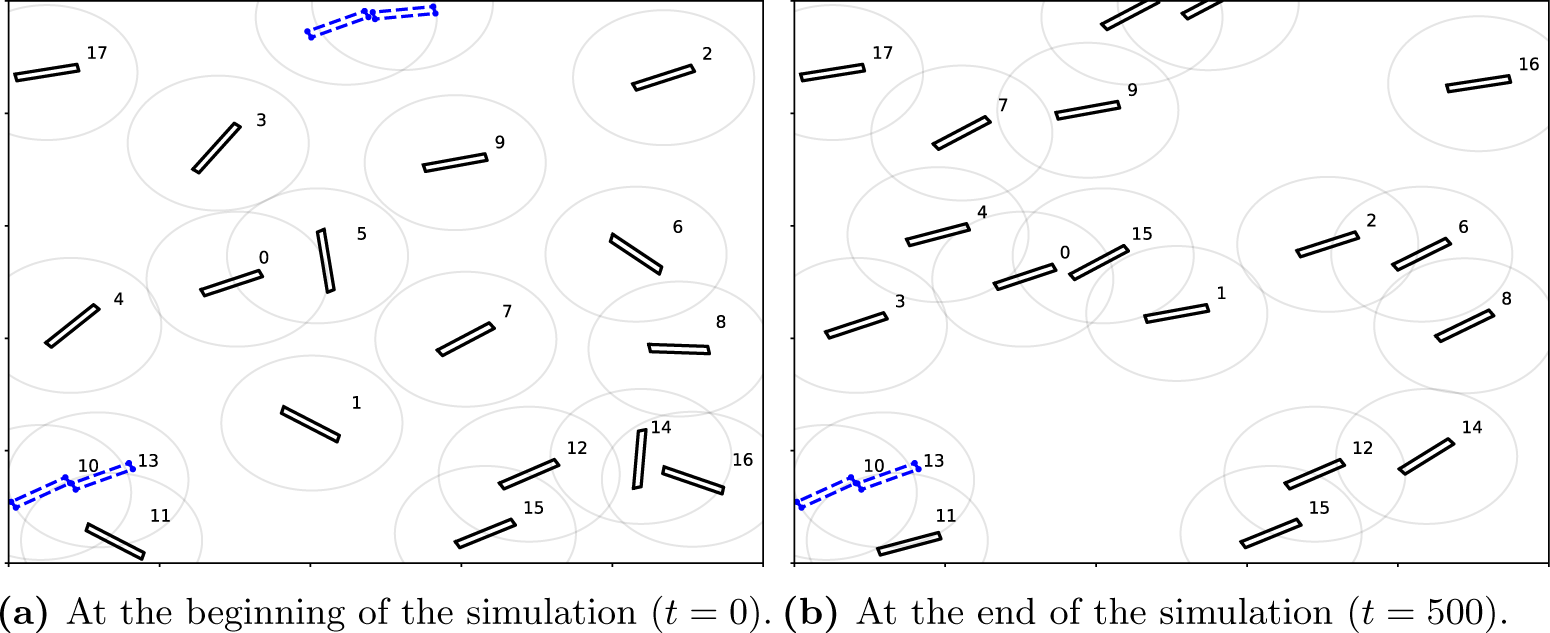
A screenshot of the simulation. Blue-dashed agents collided in the current simulation step.

### 4.3 Baseline

We used the obtained VCMCA model to evaluate the CM, CA, and CMCA scores over time. We define the scores as the complement values to the objective functions of CM (Eq. 6), CA (Eq. 7), and CMCA (Eq. 8). Thus, higher values of CM or CA scores signify better flocking performance or lower collision rates, respectively. We compared these results to a model obtained in the same way but with *ζ* = 1 rather than *ζ* = 0.5. Namely, in this configuration, the CEL algorithm aims to optimize the CM while ignoring CA. We refer to this model as CM model. In addition, we compare the proposed model with a random walk. The random walk model assumes a uniform distribution probability for each action and picks one action each time at random.

The results of the CM, CA, and CMCA scores are shown as mean ± standard deviation of *n* = 30 simulations in Figs. 6a, 6b, and 6c, respectively.

**Fig 6.**
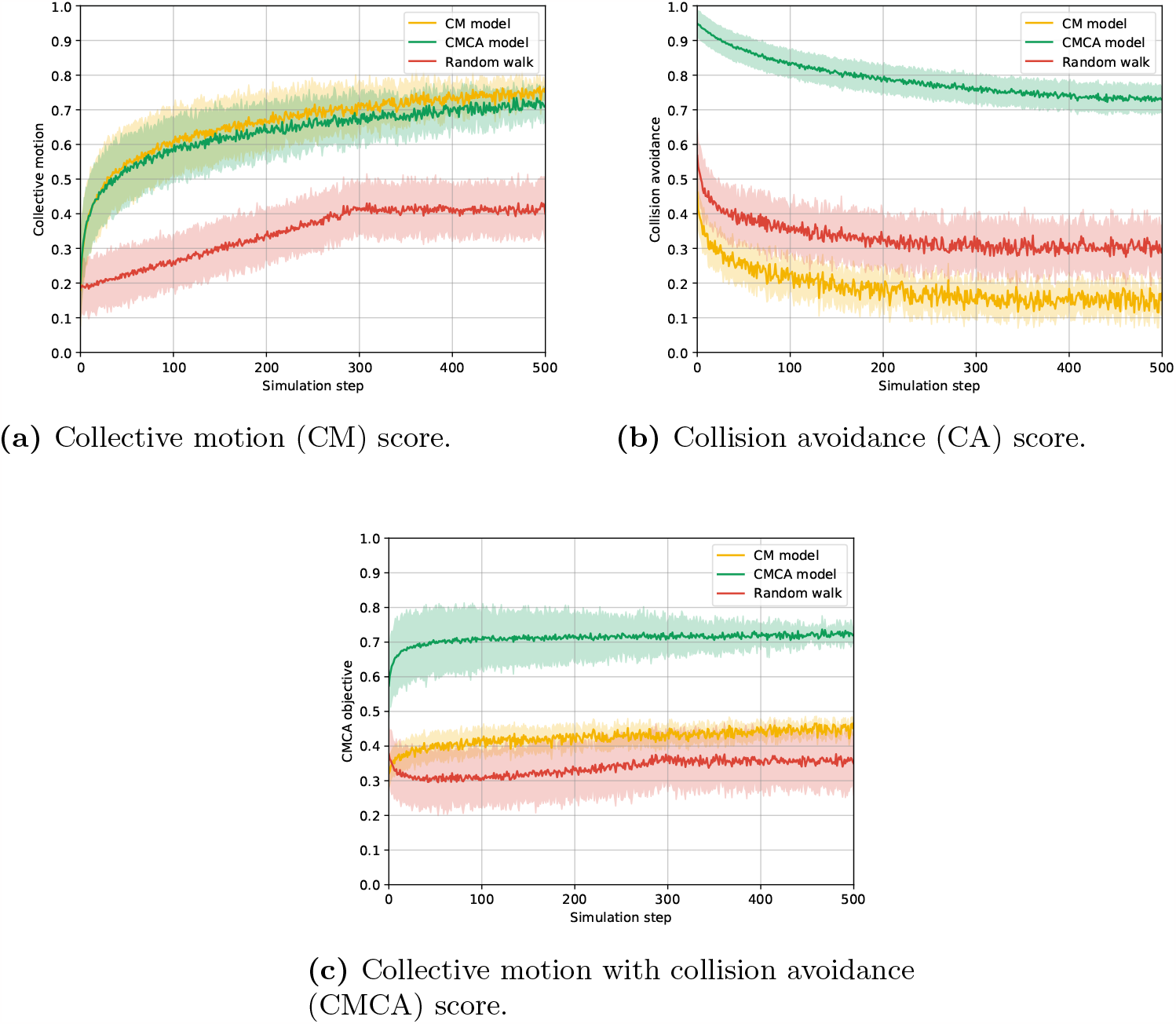
The collective motion, collision avoidance, and CMCA score of the VCMCA algorithm, VCMCA algorithm without CA, and random walk over the simulation steps for *n* = 30 repetitions. Namely, the error bars represent one standard deviation. The models’ scores are defined as the complement values of the respective objective functions.

As expected, both the CM and VCMCA models obtain flocking with a CM score of 0.75 and 0.7 while the random walk converges to 0.4, as presented in Fig. 6a. On the other hand, while the CMCA model converges to a CA score of 0.74, the random walk and CM models obtained 0.29 and 0.17, respectively, as shown in Fig. 6b. Of note, the CM model obtains the worse CA score out of the three. Since the CM model being trained only to flock with other agents, it disregards the collisions in its training, causing the agent to move closer to each other as shown in [48].

In addition, the mean results of each model over time are shown in Table 2 such that the results are the mean ± standard deviation computed on the values presented in Figs. 6a, 6b, and 6c, respectively. As seen in Fig. 6a, the agents’ convergence to cohesive movement, using the CM and CMCA models, emerges gradually with the collective motion index increasing with slowing rates over time. This process is similar to displays of hopper-bands formations, in which desert locust nymphs gather together into flocks of thousands of individuals. Due to the initially disordered state, it takes time for the visual information to be mediated between the swarm members and for them to change and adapt their heading on an individual level, which later results in the swarm reaching a coordinated heading direction on a group level. Another notable aspect is the collision rate (Fig. 6b) for CM and Random models reaching its final value in a relatively short period, in contrast to the CMCA model that decreases moderately over the time of the simulation, reaching a saturation value more than four times larger than the CM model. The gradual decrease in the CA score (that is by definition equal to an increase in the collision rate) in the CMCA model could be attributed to the CM component causing the agents to align and follow each other, in turn bringing them closer together. These results are aligning with the biological inspiration since flock members facilitate coherent movement partly for their collision-avoidance abilities [69, 70], preventing them from constantly colliding into each other, which could potentially cause high mortality in flocking animals. The models’ mean scores for each evalutaion metric are summarized in Table 2.

**Table 2.**
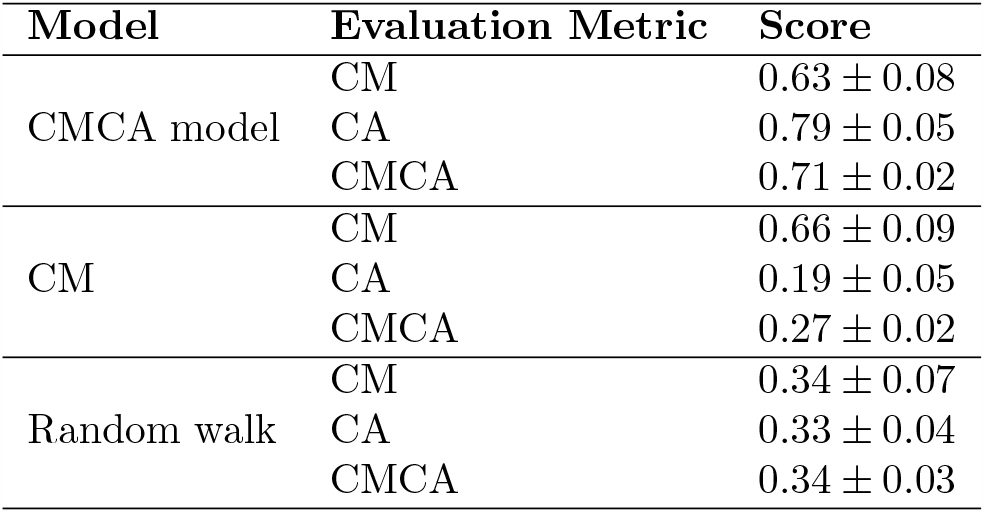
A mean ± standard deviation of the CMCA, CM, and CA scores for the CMCA model, CM model, and random walk for *n* = 30 repetitions.

### 4.4 Parameter sensitively analysis

To estimate the sensitivity of the population density, agent’s height to width ratio, and sensing radius on the VCMCA model’s performance. Given the trained VCMCA model from the CEL algorithm, we computed the CMCA score over *n* = 30 simulations with different population sizes, as presented in Fig. 7. The results are shown as mean ± standard deviation. Notably, the results are obtained after the VCMCA model is obtained on the default values (see Table 1) and evaluated on the other configurations. The results indicate an optimal flock size (*P* = 40), suggesting a strong link between the population’s size and flocking emergence. For small population sizes, the density is insufficient for phase transformation to occur since interactions between the moving agents are scarce and happen across large time intervals, thus preventing the ’directional cluster’ aggregation. In other words, agents’ behavior is similar to random-walk since local interactions do not translate to the group level. An increase in population size allows for the accumulation of aligning inter-agent interactions, in which a small cluster of agents’ incoherent movement gradually aligns the rest of the swarm resulting in a large-scale coordinated motion reminiscent of natural flocks. Further increasing the population size leads to an inherent raise in the collision rate or CA maneuvers, where agents are steering away from an aligned direction to avoid an imminent collision with an approaching neighbor. Since our model strives to minimize the collision rate, relatively large populations hinder the aligning mechanism since the CA mechanism dominates the CM mechanism. Moreover, as the size of the environment was kept constant in all the experiments, there is a linear correlation between the population’s size and density: *σ ≡* | *P* | /(*W · H*), where *σ* is the population’s density. Hence, one can obtain the same connection between the population’s (swarm) density and the flocking emergence.

**Fig 7.**
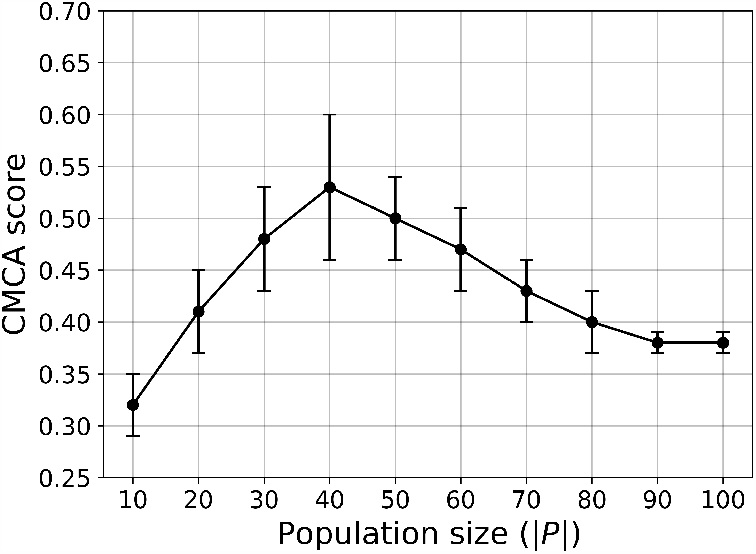
A mean ± standard deviation of the CMCA score (Eq. (8)) as a function of the population size (| *P*|) for *n* = 30 repetitions. Namely, the error bars represent one standard deviation

In a similar manner, we computed the influence of the agents’ height to width ratio (e.g. *h/s*) on the CMCA score over *n* = 30 simulations, as shown in Fig. 8.

**Fig 8.**
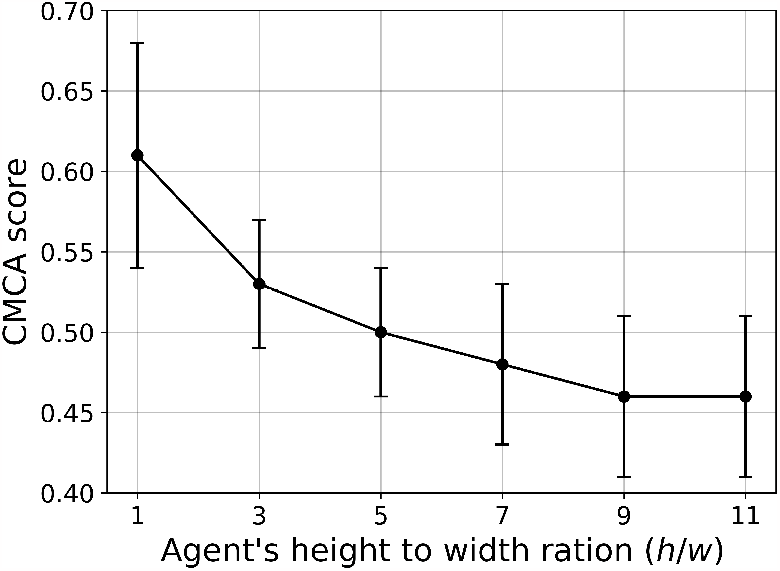
A mean ± standard deviation of the CMCA score (Eq. (8)) as a function of the agent’s ratio between the height and the width (*h/w*) for *n* = 30 repetitions Namely, the error bars represent one standard deviation.

Expectedly, larger height-to-width ratios yield lower CMCA scores. As mentioned in Section 2, elongated agent morphologies, as opposed to circular, complicate the processing of visual stimuli. Neighbor’s orientations greatly affect the subtended angle *ψ*, especially in longer agents, since the same agent with the same distance to its center could have highly differing values of *ψ*. For example, observing an agent’s head or bottom vs. observing its side will greatly change the input information and introduce significant noise into the state domain. Moreover, Fig. 8 shows that square-shaped agents got the highest CMCA score. One can conclude that a square, with its similarity to a disc, can artificially improve the convergence of a vision-based model [46, 47], since it eliminates the problem of orientational noise with its one-to-one function of distance to the angular area. However, such agent morphologies are biologically inaccurate, especially when addressing flocking animals as locusts, starlings, fish, etc [71–73].

In a similar manner, we computed the influence of the agents’ sensing radius (*r*) on the CMCA score over *n* = 30 simulations, as shown in Fig. 9, such that the x-axis indicates both the sensing radius in absolute size and as the multiplier of the agent’s height.

**Fig 9.**
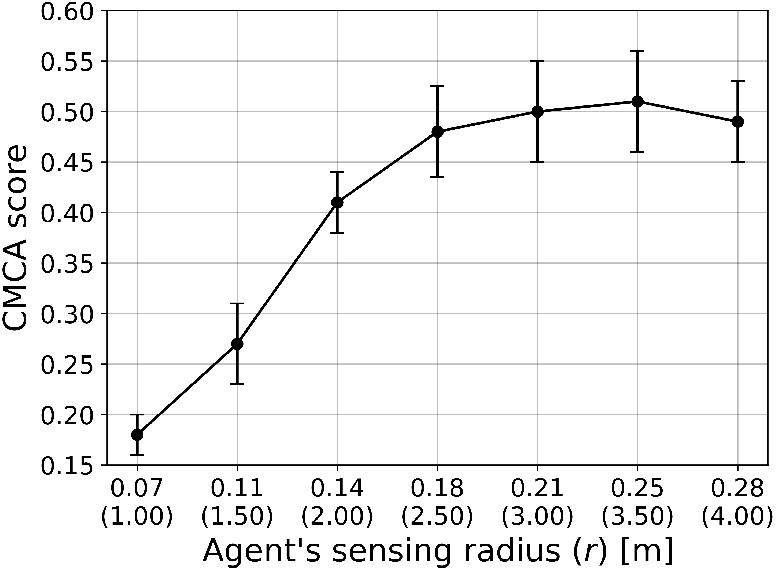
A mean ± standard deviation of the CMCA score (Eq. (8)) as a function of the agent’s sensing radius (*r*) for *n* = 30 repetitions. Namely, the error bars represent one standard deviation.

### 4.5 Explainablility of the model

The proposed model is the outcome of multiple stochastic processes, converging into collective and intelligent behavior. In this section, we aimed to measure the ability of a simple model such as Decision Tree (DT) or even its more complex but still explainable extension, the Random Forest (RF) model in capturing the complexity of the proposed model. First, we generate 1000000 cases at random, in the same way, we have done for the testing phase of the CEL algorithm (see Section 3) and query the model, resulting in a corresponding list of actions for each state. Based on this data, we train RF models with 1000 trees to allow a large number of parameters, allowing them to fit complex dynamics [64]. The obtained RF models’ accuracy relative to the VCMCA model with a restriction of the number of leaf nodes and the tree’s maximum depth is shown in Figs. 10b and 10a, respectively.

**Fig 10.**
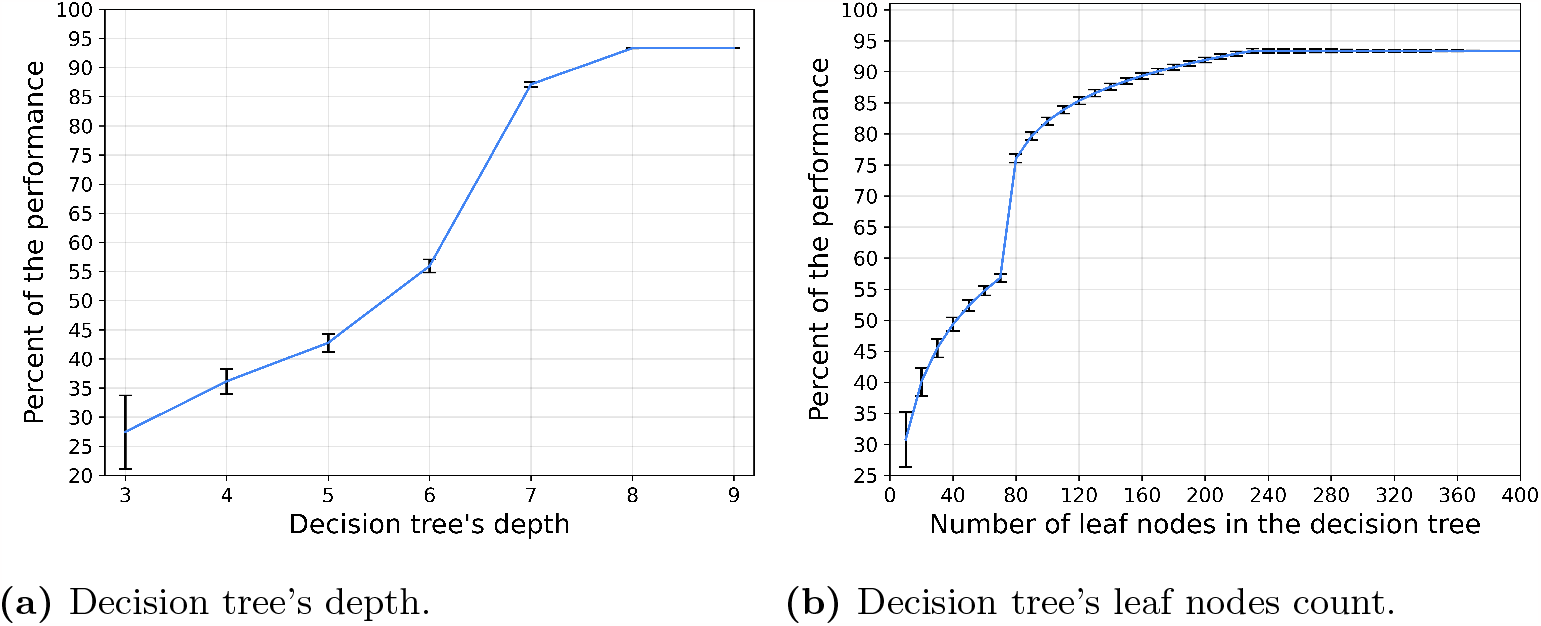
Random Forest model with 1000 trees approximated the proposed algorithm on the same set of samples. The percent of accuracy to reproduce the proposed model on the default configuration as a function of the tree’s depth and the number of leaf nodes as mean ± deviation of each decision tree is shown. Namely, the error bars represent one standard deviation.

Unsurprisingly, as more leaves are available as well as more depth to the DTs, the better the RF model approximates the VCMCA model. However, after 320 leaf nodes (or max-depth of 8) there is no significant improvement in the model’s performance over 93.37%, indicating that 6.63% of the cases are anomalies. Moreover, there is a sharp improvement between 70 leaf nodes and 80. This sharp improvement is associated with the RF model’s to differ between keeping the current vector of movement or rotating 15 degrees (e.g., the smallest turn possible in the default configuration of the model) to either the right or the left. These three actions seem to be very common and the correct separation between them causes a large performance improvement. A slight improvement can be obtained by using advance post-pruning methods such as the SAT-PP algorithm [74] but these are not significantly changing the overall observed behavior.

## 5 Discussion

In this work, we propose a vision-based collective motion with collision avoidance (VCMCA) model in a two-dimensional space. In addition, we propose a collective evolution learning (CEL) algorithm that simulates the evolution process of an individual in the context of the group (population) aiming to optimize a local VCMCA task. Individuals better perform in the task at each generation contributing to the collective knowledge, which emerges the desired multi-objective collective behavior.

We used the VCMCA model obtained from the CEL algorithm for the case of locust, showing that multi-objective swarm behavior, such as CM with CA, emerges from the optimization of the individual-level objective. These results are presented in Fig. 4 and Table 2. Moreover, Figs. 7, 8, and 9 show that the results are robust over the population’s density, agents’ shape, and sensing radius, respectively. In particular, disc-like agents with vision-based input are obtaining better results than agents with more complex topology, reproducing [46]’s results.

In addition, by approximating the obtained VCMCA model using explainable machine-learning-based models we show that small rule-based models are not able to capture the complexity of the VCMCA model and it is requiring several hundreds of rule combinations to capture only 93.3% of the VCMCA model’s ability. This model size is commonly considered out of the ability of the average person or even domain expert to explain [75]. This outcome suggests that the proposed VCMCA model obtained by the CEL is not explainable even when trying to approximate with the RF (or the DT) model, agreeing with similar results reported by [76]. Thus, while one can point to the ”reason” the RF model makes each decision, this outcome will be in the form of a majority vote between multiple boolean satisfiability problems (on the DT level), making it impractical to analyze or to extract generic roles. Nevertheless, while the proposed model is not explainable on the single decision level, the overall behavior agrees with known biological behaviors [50].

These results agree with previous outcomes proposed by [48] which shows that agents who learned secondary goals such as keeping contact with neighbors were able to keep CM. These results raise the question that if CM or CA are the primary objective of the individual, as assumed in this work, or other biological objectives such as reducing their risk of being predated or increasing their probability of finding resources are the source of these behaviors. In particular, one can evaluate the performance of CEL for *ζ* = 0 and *ζ* = 1, concluding the influence of each objective to the overall swarm behavior. In a similar manner, evaluating CEL’s performance for a topological-based rather than metric-based approach which is believed to better represent multiple scenarios in nature can shed light on the influence of the flocking mechanism on collective learning. In particular, extending the proposed model to handle three-dimensional geometry would make the model more realistic. This direction can be a promising future work.

Since the proposed VCMCA model extends the model proposed by Vicsek et al. [17], a comparison between the two is expected. Nonetheless, such a comparison is infeasible since comparing Viscak’s model as well as other similar models use different types of input data for the agents. Indeed, one would need to modify the input of these models (or algorithms) to match our own vision-based input to make the comparison legitimate. However, by doing so, the examined model would have to be altered as well to handle the new kind of input. Hence, the altered model can not be assumed to have the same behavior as the original model.

Moreover, in this work, we tackle a swarm task that assumes a group of identical agents. However, in nature, agents in the population differ by their size, capability, and even action-making process. This diversity may result in more behaviors among the agents, such as picking a leader and introducing new and complex challenges for the model.

The proposed question and many others can be addressed by introducing modifications to the proposed VCMCA model and CEL algorithm proposed in this work. As such, we believe that the proposed approach can be used as a computational background to investigate a more biologically accurate representation of swarm dynamics.

## Declarations

### Authors’ contributions

Formal analysis, investigation, and manuscript review were performed by David Krongauz; Conceptualization, formal analysis and investigation, coding, original draft preparation, and manuscript review were performed by Teddy Lazebnik.

### Code

The model’s source code is available at

https://github.com/teddy4445/Collective-Evolution-Learning.

## Acknowledgement

The research was partially funded by Chaya Rivka fund for prompting young researchers in the biomedical field.

## Appendix

### The Collective Evolution Learning algorithm time complexity

Analyzing the time and memory complexity of the proposed Collective Evolution Learning (CEL) is challenging since it uses multiple levels of stochastic processes that depended on each other. First, CEL uses the Q-learning model which converges from uniform distribution to the learned distribution up to *e*^*n*^ where *n* is the number of states in the Q-learner’s table [77]. Moreover, for each evaluation of the action’s score for the Q-learner, CEL uses the Genetic Algorithm (GA) [78]. In order to evaluate the time complexity of the GA, it can be reduced to a ”bitone” task in which the GA algorithm starts with a random population of genes represented by a string of bits and aims to make the entire population identical to a pre-defined target string of bits. The target bit string is the optimal FS represented in the same manner as the gene. As such, a binary string can be of length min_*k*_(| *S* | < 2^*k*^). While the target bit string is unavailable to us, we assume that the fitness function implicitly defines it. Thus, the GA is lower bounded by an exponent of the number of generations, *g*, and the genes’ population size, *τ*. Thus, CEL’s time complexity is *O*(*e*^*gτn*^).

### Model parameters summary

A summary of the model parameters is provided in Table 3.

**Table 3.** A summary of the parameters used in the proposed VCMCA model with their notation and detailed description.

| Parameter | Symbol | Description |
| --- | --- | --- |
| Model | $M$ | Defined by the tuple $M := (E, P)$ |
| Environment | $E$ | $E \subset \mathbb{R}^2$ |
| Population | $P$ | Fixed sized ( $\ P\ := N$ ) population of rectangular-shaped agents |
| Environment's dimensions | $H, W$ | Environment height and width, measured in agent's body-length |
| An agent | $p$ | $p \in P$, defined by a tuple $p := (l, v, a, h, w, r, \gamma)$ |
| Agent's location | $l$ | $l = (l_x, l_y) \in E$, location of the agent's center of mass in the lab frame |
| Agent's velocity | $v$ | $v \in \mathbb{R}^2$ is the current velocity vector of the agent |
| Agent's acceleration | $a$ | $a \in \mathbb{R}^2$ is the current acceleration vector of the agent |
| Agent's dimensions | $h, w$ | agent's height and width |
| Vision radius | $r$ | metric-based sensing radius of the agent, defines which agents are considered as *neighbors* |
| Vision-based input | $\gamma$ | $\gamma := (\psi, \dot{\psi}, \theta, \dot{\theta})$, parameterization of the visual input of a focal agent |
| Angular position | $\theta$ | The angle between the velocity vector of the focal agent $v_o$ and the a neighbors location $l_n$ |
| Angular area | $\psi$ | Area subtended on the retina of the focal agent |
| Local environment | $P^*$ | Subset of agents $P^*$ that are located in a sensing distance from the focal agent $P^* := \{p_i \in P : \ l_i - l_o\ < r_o\}$ |
| Visual information vector | $\nu$ | $\nu \in \mathbb{R}^{\vert \gamma\vert \cdot \vert P^*\vert }$, such that $\nu := [\gamma_1, \dots, \gamma_{\vert P^*\vert }]$ |
| The nth neighbor's vertex i | $\chi_n^i$ | The relative positions of the corners of the nth neighbor as seen by the focal agent. |
| Time -step | $t_i$ | The $i_{th}$ clock tick |
| Action | $x(t)$ | $x(t) \in \mathbb{R}^2$, Computed agent's action at time $t$ |
| CM metric | $F(P)$ | See Eq. (6) |
| Population space | $\mathbb{P}$ | The space of all possible agent populations |
| CA metric | $C(P)$ | The rate of collisions between the agents of the population to the possible number of collisions in a single point of time |
| Policy space | $\Omega$ | The space of all possible policies of the agents |
| Weight | $\zeta$ | $\zeta \in [0, 1]$ is the weight of the CM score compared to the CA score |

### Vision mechanism special cases

Following the definition of the vision mechanism described in Section 2.1, one can define three special cases for the computation of Eq. (2) following the values obtained from Eq. (3). These three cases are presented in Fig. 11.

**Fig 11.**
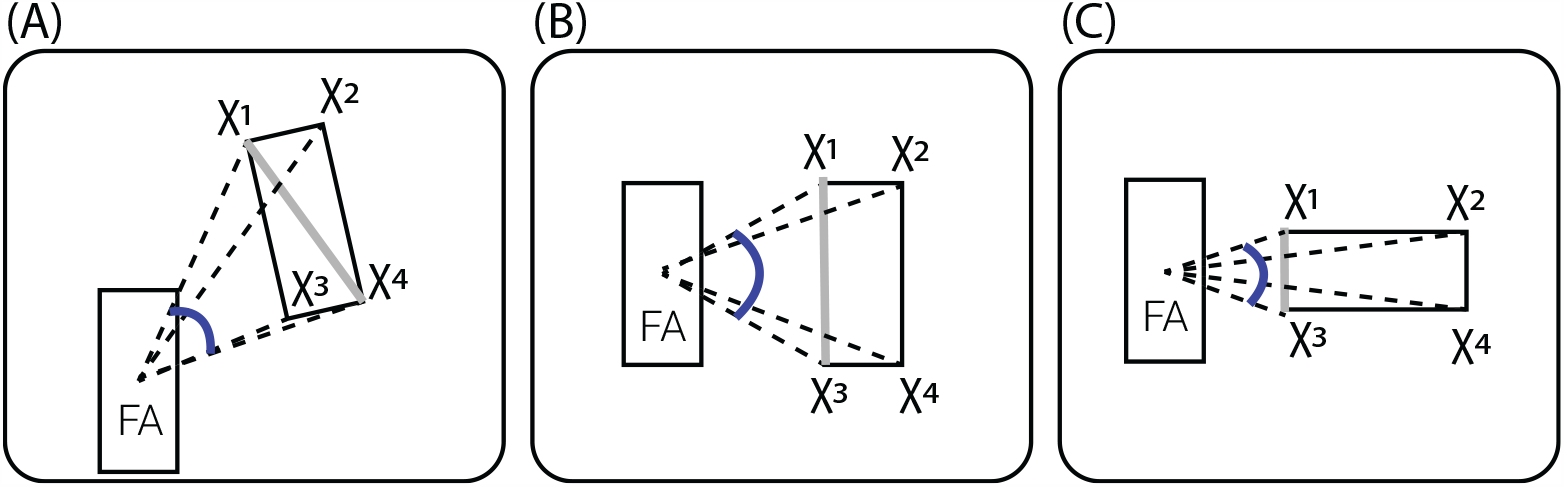
Schematic of angular area (*ψ*) calculation. FA is the focal agent and *χ*_*i*_ are the vectors connecting the FA body-center and the corners of the neighbor. The angular area *ψ*, i.e., the angle between the two extreme vectors to the neighbor, is denoted in blue. The extreme neighbor’s corners are the vertices subtending the largest possible *ψ*. This can be classified into three cases: (A) in which one of the diagonals subtends the largest angular area, or the two special cases (B, C), where the corners are the vertices on the closest neighbor’s edge to FA. The latter occurs when the line-of-sight (the vector connecting between the FA and the neighbor’s center of masses) is exactly perpendicular or parallel to the neighbor’s velocity, respectively.

